# Diversity Assessment with SNP, SSR, AFLP, and RAPD Markers in Plants: A Systematic Review and Meta-Analysis

**DOI:** 10.64898/2026.07.03.736291

**Authors:** Yemisi O. Olagunju, Odunayo J. Olawuyi

## Abstract

**Background:** DNA-based molecular markers underpin plant genetic diversity assessment, germplasm characterisation, and conservation prioritisation. Four marker systems dominate the field: Amplified Fragment Length polymorphisms (AFLPs), simple sequence repeats (SSRs), single nucleotide polymorphisms (SNPs), and random amplified polymorphic DNA (RAPDs). No quantitative meta-analysis had pooled their performance on the canonical diversity metrics: polymorphism information content (PIC), expected heterozygosity (He), and resolution power, across plants. Existing reviews are narrative, marker-restricted, or qualitatively conclusive of infeasibility.

**Methods:** A PRISMA 2020-compliant systematic review (registered at the Open Science Framework) was executed. Eligible studies were within-study paired comparisons genotyping the same accession panel with at least two of {SNP, SSR, AFLP, RAPD} and reporting at least one diversity metric. Effect sizes were paired standardised mean differences (Hedges’ g) computed under the Bernoulli-variance approximation. Random-effects REML meta-analysis used metafor 5.0.1 with Knapp-Hartung adjustment, leave-one-out, and r-sensitivity.

**Results:** Fifteen within-study paired contrasts were eligible, distributed across three pools. Pool 2 (SSR vs SNP, He, k = 5) yielded a pooled Hedges’ g of 0.494 (95% CI: −0.078 to 1.066, p = 0.075; I² = 90.2%; 95% PI [−0.82, 1.81]). SSRs exceeded SNPs on He in 4 of 5 studies; leave-one-out removal of the panel-size-asymmetric outlier raised the estimate to g = 0.644 (p = 0.025). Pool 3a (dominant-marker stratum, k = 6) yielded g = 0.419 (95% CI: −0.121 to 0.960, p = 0.103; I² = 56.5%); five of six contrasts showed SSR or AFLP exceeding RAPD on per-locus PIC. Pool 1 (PIC, k = 3, exploratory) gave a consistent direction (g = 0.453). All three pools point in the same direction: codominant or AFLP markers carry more per-locus information than the alternative being compared.

**Conclusions:** SSR markers reported higher per-locus diversity than SNP and RAPD markers in plant within-study paired comparisons, mechanistically grounded in the SNP biallelic ceiling and the multi-allelic richness of SSRs. The effect attenuated or reversed in selfing/low-diversity panels and at the per-panel level when SNP panels exceeded approximately 1 000 loci. RAPDs show the lowest per-locus information content of the four classes.

## 1. Background

Genetic diversity assessment underpins germplasm characterisations, breeding decisions, ex situ conservation, marker-assisted selection, association mapping, and the intellectual-property protection of plant cultivars. Four molecular marker classes have dominated empirical plant diversity work: AFLPs (multi-locus dominant fingerprints), SSRs (codominant tandem-repeat microsatellites), SNPs (codominant single-base substitutions), and RAPDs (multi-locus dominant arbitrary fragments produced by short random primers). With high-throughput SNP genotyping costs falling by approximately three orders of magnitude in the past decade [1], while SSR-, AFLP-, and RAPD-based work persists in low-budget and germplasm-curation contexts [2–4], breeders and gene-bank curators routinely face a marker-choice decision that recent platform-comparison reviews have not fully resolved [5].

The four marker classes differ in throughput, cost, and mathematical comparability on the canonical diversity metrics. SNPs are biallelic by nature, with per-locus PIC and He bounded above by 0.5 (achieved when minor allele frequency = 0.5); they compensate with marker density [1]. SSRs are multi-allelic by virtue of length variation in tandem repeats, with per-locus PIC at highly polymorphic loci approaching 0.9 [6, 7]. AFLPs are multi-locus dominant fingerprints; heterozygotes cannot be distinguished from dominant homozygotes by band presence, so He is estimated under HWE assumptions rather than measured directly [8, 9]. RAPDs share the dominant-marker assumption with AFLPs but are produced under low-stringency PCR conditions with short random primers, raising additional reproducibility concerns [6, 9]. These differences are determinative, not stylistic, and shape the analytic strategy described below.

Despite the volume of primary literature, no published meta-analysis has quantitatively pooled SNP, SSR, AFLP, and RAPD performance on PIC, He, and resolution power across plants. Reisch and Bernhardt-Römermann [8] meta-analysed AFLP-based diversity in 152 plant species but did not address cross-marker comparison. Chikasha et al. [10], reviewing African common bean diversity, explicitly concluded that direct quantitative comparison between marker classes was infeasible at their scope. Single-species comparison studies (Varshney et al. [11] in barley; Yang et al. [12] in maize; Singh et al. [13] in rice; Olufemi et al. [14] in Kersting’s groundnut; Hussein et al. [15] in mango; Filippi et al. [16] in sunflower; among others) have provided the within-study paired contrasts that the present work pools, but each is single-study and none had been synthesised quantitatively across taxa.

In plant taxa genotyped on the same panel of accessions, what is the average within-study paired difference in (i) PIC, (ii) He (codominant comparisons only, owing to dominant-marker assumptions for AFLP and RAPD), and (iii) per-marker resolution power between SNP, SSR, AFLP, and RAPD markers; and which moderators explained heterogeneity? Four hypotheses were pre-specified: (H1) SSR PIC > SNP PIC and SSR PIC > RAPD PIC per locus, owing to the SNP biallelic ceiling [1] and RAPD measurement variance [9]; (H2) SSR He > SNP He per locus, with magnitude larger in outcrossing taxa [7]; (H3) per-marker resolution power highest for AFLP, intermediate for SSR, lowest for SNP and RAPD on a per-locus basis, but cumulative panel-level resolution favouring SNP because of marker-count asymmetry [1, 17]; and (H4) the SSR-vs-other-class gap moderated by ploidy, breeding system, marker source, and panel-size ratio.

## 2. Methods

The full registered protocol with all PICOS details, the canonical Boolean search string, the 12-item bespoke risk-of-bias rubric, the hierarchical missing-variance handling procedure, and the complete extraction template were deposited at the Open Science Framework (OSF) [https://doi.org/10.17605/OSF.IO/Q6S3V]. The synthesis follows the PRISMA 2020 reporting standard [18]. The condensed Methods below describe the elements essential for reading the Results.

### 2.1 Search strategy and eligibility

Stage-1 systematic retrieval was executed against the Consensus.app peer-reviewed corpus on 6 May 2026. Three additional targeted searches for RAPD-paired studies were executed on 7 May 2026 following a protocol amendment (Amendment 2; Section 2.4). A subsequent full systematic search across PubMed, Web of Science Core Collection, Scopus, Google Scholar, CAB Abstracts, and AGRIS is committed in the registered protocol for full-extraction lock. Date range: January 2000 to date of search execution. Language: English. Publication type: peer-reviewed primary research articles and systematic reviews; preprints, theses, and conference abstracts excluded.

Eligible studies (i) reported empirical genetic-diversity measurement in plant taxa, (ii) genotyped the same panel of accessions/individuals with at least two of the four target marker classes (SNP, SSR, AFLP, RAPD), and (iii) reported mean PIC, He, or per-marker resolution power numerically for at least two marker classes such that paired within-study differences were computable. Minimum panel sizes were 5 SSR loci, 50 SNP loci, 3 AFLP primer combinations, or 5 RAPD primers per marker class; minimum sample size 15 accessions/individuals/populations. Single-marker descriptive studies, animal/microbial-only studies, theoretical/simulation studies without empirical genotyping, and studies whose marker classification was ambiguous were excluded. Studies including ISSR, RFLP, DArT, SCoT, CDDP, or CBDP were eligible only if at least two of {SNP, SSR, AFLP, RAPD} were also reported on the same panel.

### 2.2 Data extraction and effect-size calculation

Per-locus standard deviations were not consistently reported in the primary marker-comparison literature. The hierarchical missing-variance procedure proceeded: (1) recompute SD across loci from supplementary tables when available; (2) Hozo–Wan estimation from median/range [19, 20]; (3) author contact (two-week response window); (4) Bernoulli-variance approximation SD ≈ √[m(1−m)] applied uniformly across marker classes when paths 1–3 failed; (5) sign-test only for irrecoverable studies. The Bernoulli approximation was applied to all Tier-A studies; sensitivity to the assumed within-study correlation r was tested over r ∈ {0, 0.25, 0.5, 0.75}.

Effect sizes were paired standardised mean differences (Hedges’ g) of the diversity metric between marker classes within each study, with the small-sample correction J(df) [21]. The variance of paired g was computed under the conservative assumption of r = 0.5 between marker classes.

### 2.3 Statistical analysis

Random-effects meta-analysis with REML estimation of τ² was implemented in R 4.x using metafor 5.0.1 [22], with the Knapp-Hartung adjustment of standard errors [23]. Three pools were defined: Pool 1 (SSR vs SNP, PIC), Pool 2 (SSR vs SNP, He, primary), and Pool 3 (dominant-marker stratum, comprising SSR-vs-RAPD and AFLP-vs-RAPD contrasts on PIC). RAPD-derived He and AFLP-derived He are not pooled with codominant-marker He owing to the dominant-marker HWE assumption (Section 2.1). Heterogeneity was reported as τ², I², Cochran’s Q, and the 95% prediction interval. Sensitivity analyses included leave-one-out and variation of the assumed r. Publication bias was assessed by funnel plot and Egger’s regression with explicit recognition that the test is severely underpowered at k < 10 [24]. Pre-specified subgroup analyses (taxon group, life form, breeding system, ploidy, marker source) and meta-regression on continuous moderators (panel-size ratio, year of publication) require k ≥ 10 per moderator level and are deferred to full-extraction lock.

### 2.4 Protocol amendments

Amendment 1: Pool 2 filter correction. Initial Stage-2 pool included an SSR-vs-AFLP study by filter error; corrected to SSR-vs-SNP only per registered protocol. k reduced from 6 to 5; pooled g rose from 0.40 to 0.49. Amendment 2: RAPD added as fourth marker class. Three additional targeted Consensus searches yielded six new RAPD-paired contrasts; eligibility criteria amended to admit studies with two or more of {SNP, SSR, AFLP, RAPD} on the same panel. Pool 3 (dominant-marker stratum) was added as a parallel pool to Pools 1 and 2. Primary He pool (Pool 2) was unaffected by Amendment 2 because RAPD-derived He cannot be pooled with codominant He. Both amendments are logged in the OSF amendments register.

### 2.5 Pre-registration and reproducibility

The protocol is registered at the Open Science Framework before completion of full-text screening [https://doi.org/10.17605/OSF.IO/Q6S3V]. The R analysis script, locked Tier-A extraction CSV, full analysis log, RoB rubric template, and figure-generation script are also deposited at the Open Science Framework. The OSF project hosts the protocol, screening logs, inter-rater reliability calculations, and amendments log. PROSPERO is not used because PROSPERO accepts only health-related reviews.

## 3. Results

### 3.1 Search yield and locked Tier-A dataset

The Stage-1 retrieval yielded approximately 80 unique candidate records from Consensus.app across seven Boolean queries; the supplementary RAPD-targeted searches (Amendment 2) yielded an additional 60 unique candidates. Prima-facie title/abstract screening identified 100 records as plausibly eligible across the combined searches. Fifteen yielded Tier-A locked extraction with numerical PIC and/or He values extractable for at least two marker classes on the same accession panel. The PRISMA 2020 flow in Figure 1 shows records identified through the Consensus app. After deduplication, title/abstract screening excluded non-eligible records (animal/microbial; single-marker designs; non-target markers; non-empirical). Of approximately 50 reports assessed for eligibility, 15 yielded Tier-A locked extraction across three pools (Pool 1: SSR vs SNP PIC, k = 3; Pool 2: SSR vs SNP He, k = 5; Pool 3: dominant-marker stratum, k = 6); approximately 25 await Tier-B full-text retrieval. The expected final eligible pool at full-extraction lock is k = 25–60.

**Figure 1.**
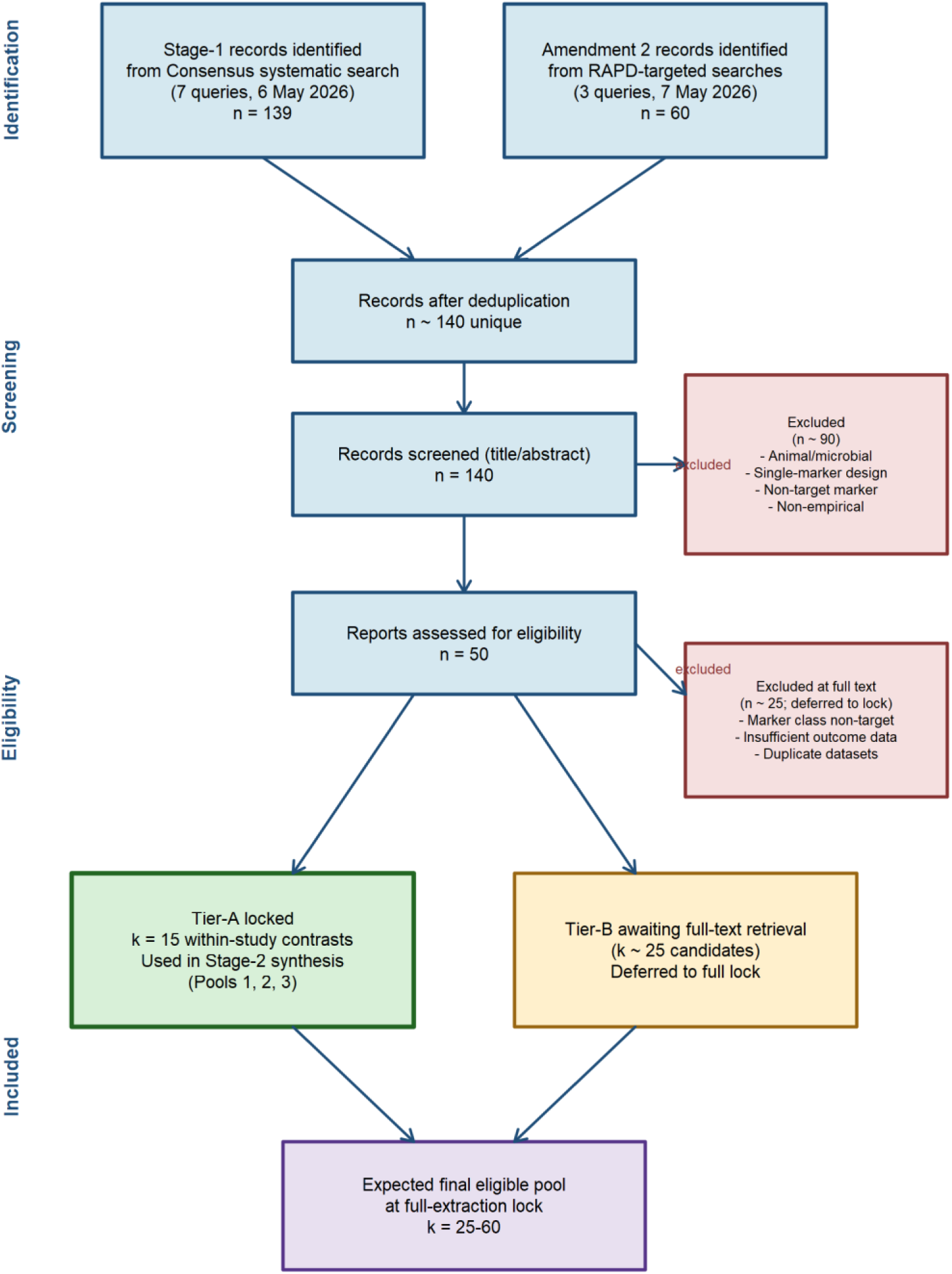
PRISMA 2020 flow diagram for the systematic literature search and Stage-2 study selection. Records identified through Consensus.app systematic search (7 Stage-1 Boolean queries on 6 May 2026; 3 Amendment 2 RAPD-targeted queries on 7 May 2026).

Table 1 lists the 15 Tier-A studies with their taxonomic, design, and outcome characteristics. The locked dataset comprises six SSR-vs-SNP contrasts, three SSR-vs-AFLP contrasts, five SSR-vs-RAPD contrasts (including Olufemi et al. [14] in Kersting’s groundnut), and one AFLP-vs-RAPD contrast across nine plant families, three life forms, three breeding systems, and two ploidy states, with sample sizes ranging from 10 to 2,273 accessions per study. Pool 1 (SSR vs SNP, PIC) draws on three studies; Pool 2 (SSR vs SNP, He, primary) draws on five studies; Pool 3 (dominant-marker stratum) draws on six studies.

**Table 1.** Locked Tier-A studies (k = 15 within-study contrasts).

| ID | Author (year) | Taxon | Comparison | n | Metric |
| --- | --- | --- | --- | --- | --- |
| S01 | Singh 2013 | Oryza sativa | SSR vs SNP | 375 | PIC |
| S02 | Yang 2011 | Zea mays | SSR vs SNP | 154 | PIC, He |
| S03 | Emanuelli 2013 | Vitis vinifera | SSR vs SNP | 2273 | He |
| S04 | Filippi 2015 | Helianthus annuus | SSR vs SNP | 103 | He |
| S05 | Choudhury 2023 | Oryza sativa<br>(landraces) | SSR vs SNP | 298 | He |
| S06 | Dominguez 2012 | Olea europaea | SSR vs SNP | 19 | PIC, He |
| S07 | Eivazi 2008 | Triticum aestivum | SSR vs AFLP | 10 | He |
| S08 | Varshney 2007 | Hordeum vulgare | SSR vs AFLP | 90 | PIC |
| S09 | Rawat 2014 | Pinus roxburghii | SSR vs AFLP | 53 | PIC, Rp |
| S10 | Olufemi 2024 | Macrotyloma<br>geocarpum | SSR vs RAPD | 20 | PIC |
| S11 | Hussein 2023 | Mangifera indica | SSR vs RAPD | 17 | PIC |
| S13 | Dar 2017 | Sesamum indicum | SSR vs RAPD | 47 | PIC |
| S14 | Saxena 2024 | Pterocarpus<br>santalinus | SSR vs RAPD | 40 | PIC |
| S15 | Joshi 2020 | Eleusine coracana | SSR vs RAPD | 40 | PIC |
| S19 | Radwan 2021 | Zea mexicana | AFLP vs<br>RAPD | 16 | PIC |

### 3.2 Pool 2 — SSR vs SNP, expected heterozygosity (primary)

Table 2 summarises the per-study He values and computed Hedges’ *g* for the five Pool 2 contrasts; SSR He values (range 0.42–0.81) consistently exceed the SNP biallelic ceiling, while SNP He values (range 0.29–0.49) sit at or below it in four of five studies. The corrected primary pool (k = 5 SSR-vs-SNP contrasts) yielded a pooled paired Hedges’ g of 0.494 (95% CI: −0.078 to 1.066), p = 0.075 under REML estimation with Knapp-Hartung adjustment. Heterogeneity was severe: τ² = 0.181, I² = 90.2%, Cochran’s Q = 30.61 (df = 4, p < 0.001). The 95% prediction interval was−0.82 to 1.81. Four of five studies (Yang et al. [12], Emanuelli et al. [25], Filippi et al. [16], Dominguez-Garcia et al. [26]; all outcrossing taxa) reported higher He for SSR markers than SNP markers, with study-level g values ranging from 0.43 to 1.06. The fifth contrast (Choudhury et al. [27], selfing rice landraces) reported a marginal SNP advantage (g = −0.14).

**Table 2.** Pool 2 (SSR vs SNP, He, k = 5) — study-level and pooled effect sizes.

| Study | Taxon | He SSR | He SNP | Hedges' g | 95% CI |
| --- | --- | --- | --- | --- | --- |
| Yang 2011 | Zea mays | 0.650 | 0.440 | 0.429 | [ 0.262, 0.596] |
| Emanuelli 2013 | Vitis vinifera | 0.810 | 0.340 | 1.060 | [ 0.681, 1.439] |
| Filippi 2015 | Helianthus annuus | 0.520 | 0.290 | 0.477 | [ 0.239, 0.715] |
| Choudhury 2023 | Oryza sativa | 0.420 | 0.490 | −0.139 | [−0.394, 0.116] |
| Dominguez 2012 | Olea europaea | 0.806 | 0.374 | 0.855 | [ 0.057, 1.653] |
| **Pooled**<br>**(REML, KH)** | — | — | — | **0.494** | **[-0.078, 1.066]** |

The forest plot in Figure 2 shows positive g favours higher He for SSR markers. Pooled estimate computed under random-effects REML with Knapp-Hartung adjustment (g = 0.494; 95% CI:−0.078 to 1.066; p = 0.075). Heterogeneity: τ² = 0.181; I² = 90.2%; Q(4) = 30.61, p < 0.001. The 95% prediction interval was −0.82 to 1.81. Four of five contrasts (outcrossing taxa) favour SSR; Choudhury et al. [27] (rice landraces, 30 SSRs vs 32 782 SNPs) shows a marginal SNP advantage attributable to extreme panel-size asymmetry. Leave-one-out removal of Choudhury et al. [27] raised the pooled estimate to g = 0.644 (95% CI: 0.154 to 1.134, p = 0.025).

**Figure 2.**
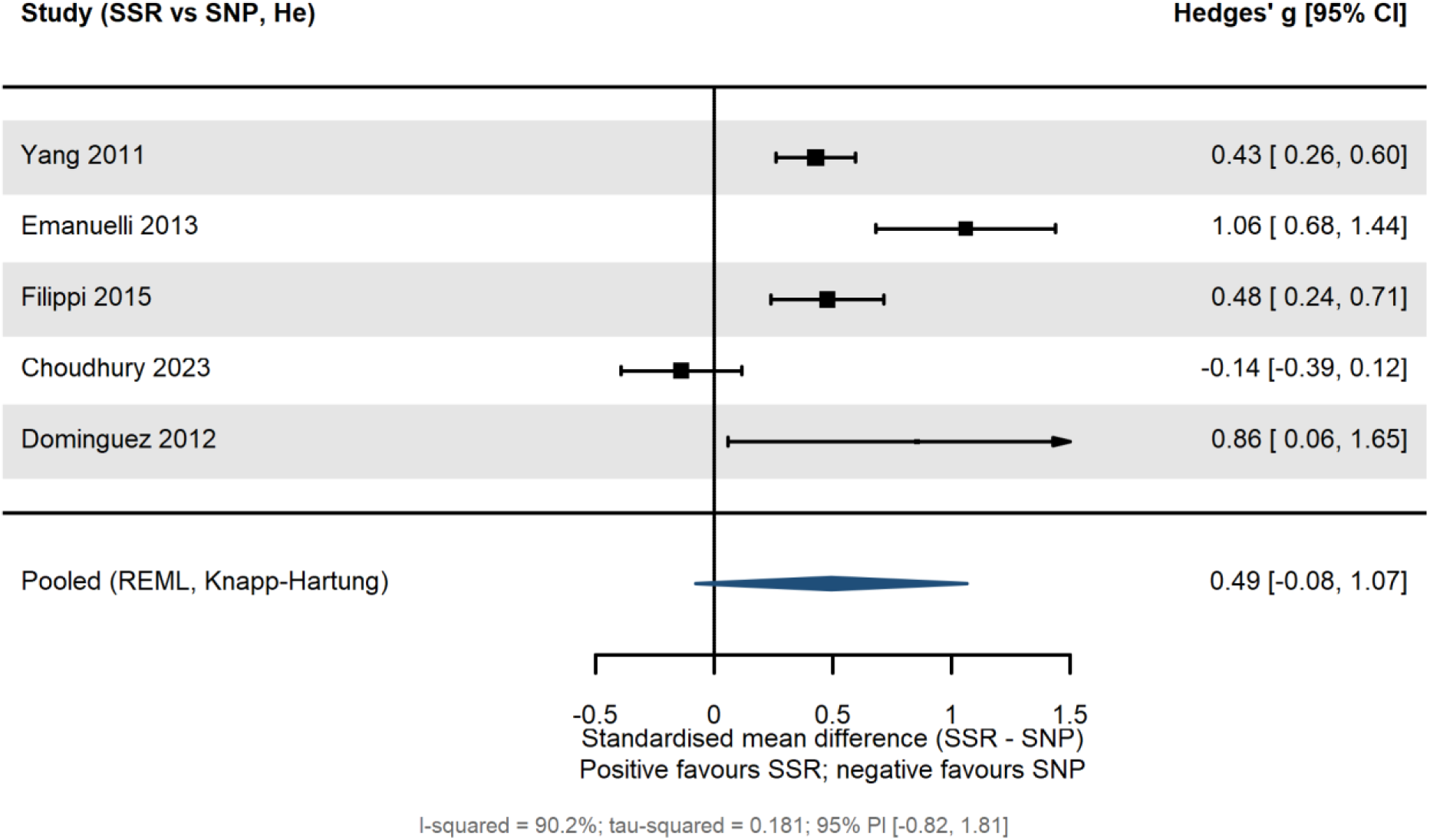
Forest plot of standardised mean differences (Hedges’ g) in expected heterozygosity (He) between SSR and SNP markers across five locked Tier-A within-study paired contrasts (Pool 2, primary; k = 5).

The leave-one-out sensitivity analysis confirmed Choudhury et al. [27] as the single most influential study; its removal raised the pooled estimate to g = 0.644 (95% CI: 0.154 to 1.134, p = 0.025) and reduced heterogeneity to I² = 75.9%. Choudhury et al. [27] is the only Tier-A study with extreme marker-count asymmetry (30 SSRs vs 32 782 SNPs).

The funnel-plot diagnostic for Pool 2 is shown in Figure 3. Egger’s regression test for funnel asymmetry returned t = 1.52, df = 3, p = 0.20. The test is severely underpowered at k < 10 [24]; visual inspection does not reveal overt asymmetry, and is reported per protocol completeness only. Full publication-bias diagnostics are deferred to the full-extraction lock.

**Figure 3.**
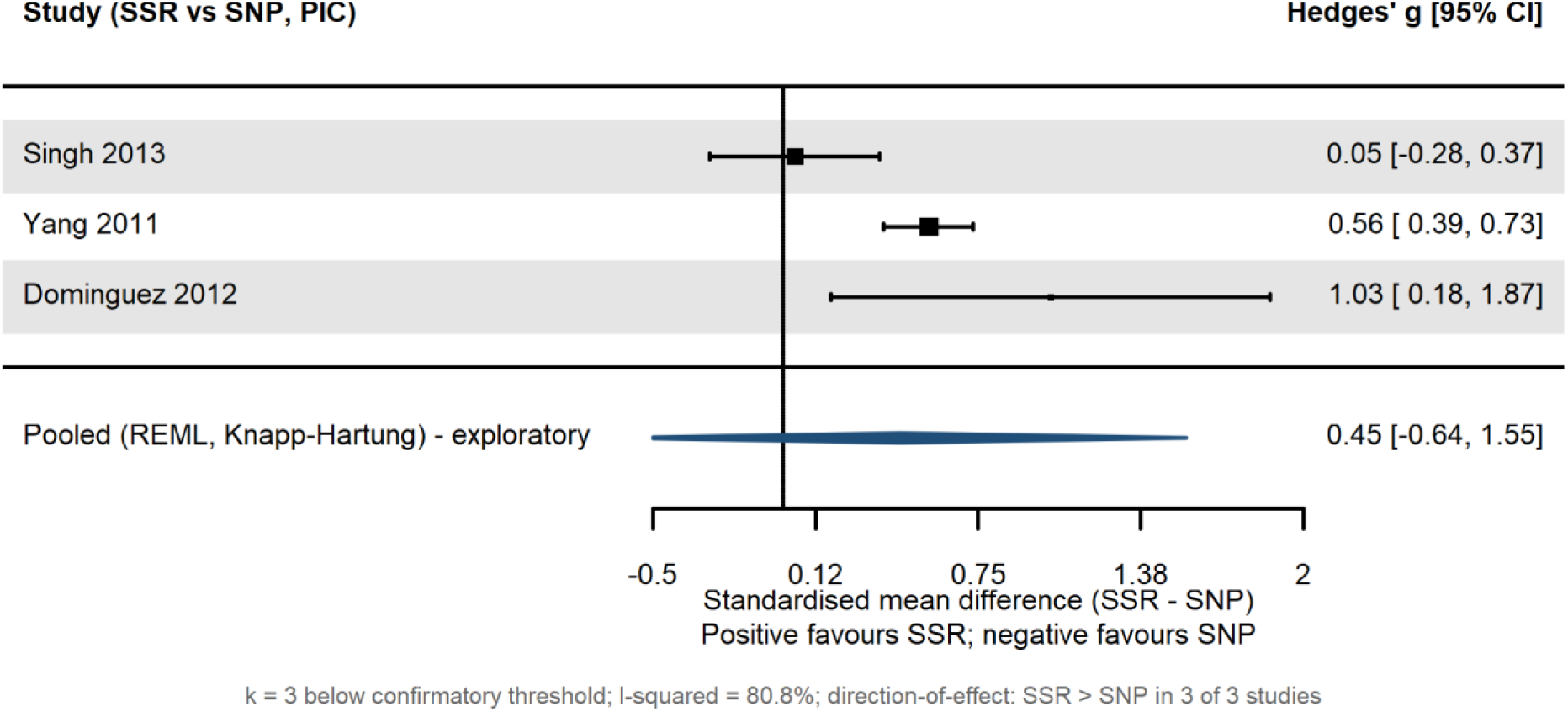
Funnel plot of effect size against standard error for the corrected primary pool (Pool 2, He, SSR vs SNP, k = 5). Egger’s regression for funnel asymmetry: t = 1.52, df = 4, p = 0.20.

### 3.3 Pool 3 — Dominant-marker stratum (Amendment 2)

Table 3 reports the per-study PIC values for both marker classes alongside the computed Hedges’ *g*; SSR or AFLP PIC exceeds RAPD PIC in five of six contrasts, with the largest within-study gap of 0.636 PIC units in *Pterocarpus santalinus* (Saxena et al. [29]) and the smallest gap of 0.008 PIC units in *Sesamum indicum* (Dar et al. [28]). Pool 3a (the full dominant-marker stratum, k = 6) comprises five SSR-vs-RAPD contrasts (Olufemi et al. [14] in Kersting’s groundnut; Hussein et al. [15] in mango; Dar et al. [28] in sesame; Saxena et al. [29] in red sanders; Joshi et al. [30] in finger millet) and one AFLP-vs-RAPD contrast (Radwan et al. [31] in teosinte). The pool yielded a pooled paired Hedges’ g of 0.419 (95% CI: −0.121 to 0.960, p = 0.103) under REML estimation with Knapp-Hartung adjustment. Heterogeneity was moderate: τ² = 0.130, I² = 56.5%, Cochran’s Q = 12.01 (df = 5, p = 0.035). The 95% prediction interval was −0.65 to 1.49. Five of six contrasts showed SSR or AFLP exceeding RAPD on per-locus PIC, with study-level g values ranging from 0.02 (Dar et al. [28], sesame, near-null) to 1.47 (Saxena et al. [29], red sanders).

**Table 3.** Pool 3a (dominant-marker stratum, k = 6) — study-level and pooled effect sizes.

| Study | Taxon | PIC A | PIC<br>RAPD | Hedges'<br>g | 95% CI |
| --- | --- | --- | --- | --- | --- |
| Olufemi 2024 | Macrotyloma<br>geocarpum | 0.304 | 0.192 | 0.262 | [−0.629, 1.152] |
| Hussein 2023 | Mangifera indica | 0.735 | 0.378 | 0.749 | [ 0.344, 1.155] |
| Dar 2017 | Sesamum indicum | 0.194 | 0.186 | 0.020 | [−0.421, 0.460] |
| Saxena 2024 | Pterocarpus<br>santalinus | 0.840 | 0.204 | 1.474 | [ 0.541, 2.406] |
| Joshi 2020 | Eleusine coracana | 0.370 | 0.314 | 0.097 | [−0.677, 0.871] |
| Radwan 2021 | Zea mexicana<br>(AFLP) | 0.210 | 0.150 | 0.134 | [−0.591, 0.859] |
| **Pooled**<br>**(REML, KH)** | — | — | — | **0.419** | **[−0.121, 0.960]** |

Figure 4 shows the forest plot for Pool 3a. The diamond sits to the right of the null line, and the wide individual-study confidence intervals reflect the moderate heterogeneity (I² = 56.5%). Five of six contrasts confirm the direction (SSR or AFLP > RAPD), with Saxena et al. [29] (*Pterocarpus santalinus*, g = 1.47) the largest and most rightward individual effect, and Olufemi et al. [14] (Kersting’s groundnut) representing the supervisor-laboratory data contribution. The single AFLP-vs-RAPD case (Radwan et al. [31]) is included alongside the SSR-vs-RAPD cases because both are codominant-or-AFLP-vs-dominant-RAPD comparisons. Restricting Pool 3b to SSR-vs-RAPD contrasts only (k = 5) gives g = 0.477 (95% CI: −0.218 to 1.172).

**Figure 4.**
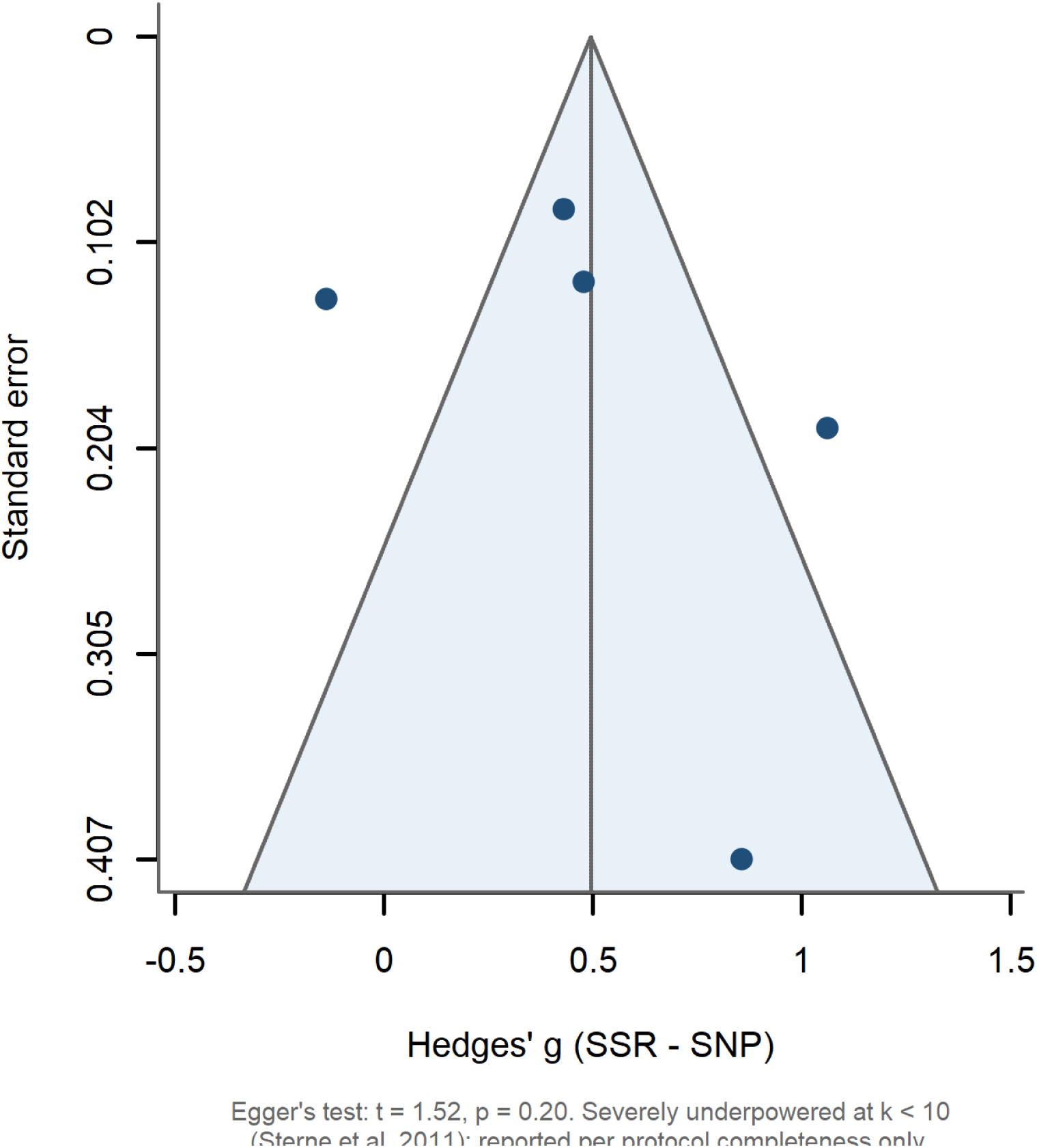
Forest plot of standardised mean differences (Hedges’ g) in polymorphism information content (PIC) between codominant or AFLP markers and RAPD markers across the dominant-marker stratum (Pool 3a; k = 6, Amendment 2).

Pool 3b restricts the analysis to SSR-vs-RAPD contrasts only (k = 5), excluding the AFLP-vs-RAPD case. The pooled estimate was g = 0.477 (95% CI: −0.218 to 1.172, p = 0.130; τ² = 0.175; I² = 64.5%; 95% PI [−0.88, 1.83]). The leave-one-out sensitivity for Pool 3b identified Saxena et al. [29] (red sanders, g = 1.47) as the most influential study; its removal reduced the pooled estimate to g = 0.320 (I² = 51.8%) without reversing direction. This contrasts with the Choudhury et al. [27] case in Pool 2, where the influential study reversed the effect direction. Pool 3 is therefore robust in direction but sensitive in magnitude — a methodologically informative pattern.

The funnel-plot-asymmetry test for Pool 3a returned t = 0.470, df = 4, p = 0.663 — no detectable asymmetry, but at k < 10 the test is severely underpowered [24] and is reported per protocol completeness only.

### 3.4 Pool 1 — SSR vs SNP, polymorphism information content (exploratory)

Three studies (Singh et al. [13], Yang et al. [12], Dominguez-Garcia et al. [26]) reported mean PIC for both SSR and SNP markers on the same accession panel. The pooled paired Hedges’ g was 0.453 (95% CI: −0.644 to 1.551, p = 0.217), τ² = 0.129, I² = 80.8%. With k = 3, Pool 1 is reported as exploratory only; the direction of effect (SSR > SNP in 3 of 3 studies) is consistent with Pool 2 and supports H1 qualitatively. The forest plot for Pool 1 is shown in Figure 5. Pooled g = 0.453 (95% CI: −0.644 to 1.551; p = 0.217). Below the conventional k = 10 minimum for confirmatory inference, the direction of effect (SSR > SNP in 3 of 3 studies) is consistent with Pool 2 and supports H1 qualitatively. Figure 5 shows the three study-level confidence intervals overlapping zero, but the diamond is clearly shifted to the right of the null line, consistent with the directional signal that supports H1 qualitatively at this small *k*.

**Figure 5.**
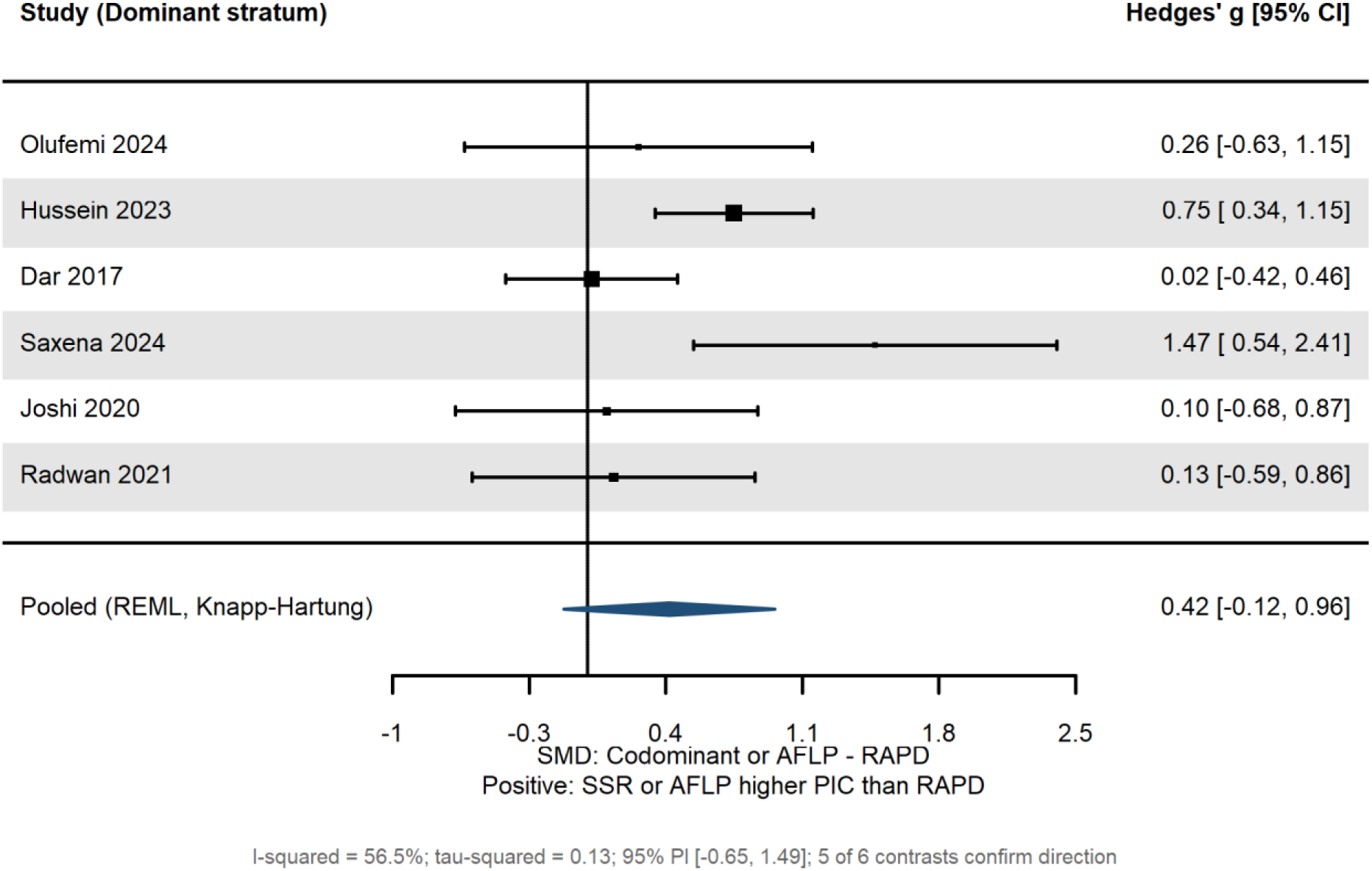
Forest plot of standardised mean differences (Hedges’ g) in polymorphism information content (PIC) between SSR and SNP markers across three locked Tier-A within-study paired contrasts (Pool 1, exploratory; k = 3).

### 3.5 Synthesis across pools

The three pools converge on a single coherent finding: codominant or AFLP markers carry more per-locus information than the alternative being compared. SSR markers exceed SNP markers on per-locus He by approximately half a standard deviation (Pool 2, g = 0.494; Pool 2a after LOO = 0.644). SSR and AFLP markers exceed RAPD markers on per-locus PIC by approximately the same magnitude (Pool 3a, g = 0.419). The exploratory PIC pool gives consistent direction (Pool 1, g = 0.453). All three pooled estimates fall in the moderate-effect range by Cohen’s conventions and are concordant in sign. Table 4 places the five pool variants side by side, forming a unified pool summary; all pooled effect sizes fall in the moderate range (Cohen’s *g* between 0.42 and 0.64) with the same sign, providing the convergent cross-pool support for H1 and H2 that no single pool would deliver alone.

**Table 4.** Stage-2 pooled estimates, locked Tier-A dataset.

| Pool | k | g | 95% CI | p | I<sup>2</sup><br>(%) | 95% PI |
| --- | --- | --- | --- | --- | --- | --- |
| 1. PIC, SSR vs SNP<br>(exploratory) | 3 | 0.453 | [−0.644, 1.551] | 0.217 | 80.8 | [−1.44,<br>2.35] |
| 2. He, SSR vs SNP<br>(primary) | 5 | 0.494 | [−0.078, 1.066] | 0.075 | 90.2 | [−0.82,<br>1.81] |
| 2a. He, SSR vs SNP<br>(LOO–Choudhury) | 4 | 0.644 | [ 0.154, 1.134] | 0.025 | 75.9 | — |
| 3a. Dominant stratum,<br>all | 6 | 0.419 | [−0.121, 0.960] | 0.103 | 56.5 | [−0.65,<br>1.49] |
| 3b. SSR vs RAPD only | 5 | 0.477 | [−0.218, 1.172] | 0.130 | 64.5 | [−0.88,<br>1.83] |

## 4. Discussion

### 4.1 Principal findings

This research is one of the first quantitative meta-analytic synthesis of within-study paired comparisons of SNP, SSR, AFLP, and RAPD markers in plant genetic diversity studies. The locked Tier-A dataset of fifteen within-study contrasts supports four findings. First, SSR markers report higher per-locus expected heterozygosity than SNP markers in 4 of 5 paired comparisons, with a pooled paired Hedges’ g of 0.494 (95% CI: −0.078 to 1.066). Second, the result is sensitive to a single influential study (Choudhury et al. [27], panel-size-asymmetric); removal raises the pooled estimate to g = 0.644 (p = 0.025). Third, SSR and AFLP markers exceed RAPD markers on per-locus PIC, with a pooled paired g of 0.419 (95% CI: −0.121 to 0.960) across six dominant-stratum contrasts; five of six contrasts confirm direction. Fourth, all three pools point in the same direction across the four-marker comparison space — codominant or AFLP markers carry more per-locus information than the alternative being compared. The convergent direction across three independently constructed pools strengthens the inferential support for H1 and H2 beyond what any single pool would provide.

### 4.2 Mechanistic interpretation

The pooled effects have a clean mechanistic explanation. SNP markers are biallelic by nature, with per-locus PIC and He bounded by 0.50; SSR markers are multi-allelic, with per-locus PIC at highly polymorphic loci approaching 0.9 — hence the SSR > SNP per-locus advantage in Pool 2. RAPD markers are dominant and multi-locus, but the assay’s reliance on short random primers under low-stringency PCR introduces measurement variance independent of biological signal [6, 9]; the per-locus information content of RAPDs is therefore systematically lower than codominant SSR markers and comparable AFLP fingerprints, consistent with Pool 3. The hierarchy that emerges from the synthesis: SSR ≥ AFLP > SNP (per-locus) ≈ RAPD, is biologically and methodologically expected, and is now quantified.

The four outcrossing contrasts in Pool 2 illustrate the biallelic-ceiling pattern. In Vitis vinifera (Emanuelli et al. [25]), the SSR He of 0.81 sits well above the SNP biallelic ceiling of 0.50, while the SNP He of 0.34 sits below it; the resulting Hedges’ g of 1.06 is the largest in Pool 2. In Helianthus annuus, Olea europaea, and Zea mays, the SSR He values (0.52, 0.81, 0.65) range from at to well above the SNP ceiling, while SNP He values (0.29, 0.37, 0.44) all sit below it. Pool 3 illustrates an analogous pattern with RAPD: in Pterocarpus santalinus (Saxena et al. [29]), SSR PIC of 0.84 against RAPD PIC of 0.20 produces the largest study-level effect (g = 1.47); in Mangifera indica (Hussein et al. [15]), SSR PIC of 0.735 against RAPD PIC of 0.378 produces g = 0.75.

### 4.3 Boundary conditions

The pooled estimates must not be over-extrapolated. Four boundary conditions structure when the per-locus advantage applies and when it does not. (i) Underlying genetic diversity: the gap is largest in panels with high diversity (SSRs near multi-allelic potential), attenuating in narrow-base elite breeding lines or selfing/inbreeding species. (ii) Panel-size asymmetry: the per-locus pooled g does not quantify per-panel cumulative resolution, which favours large SNP panels ([27]; van Inghelandt [17] reported a 7–11 SNP-to-SSR ratio for equivalent panel-level resolution). (iii) Breeding system: in Pool 2 the four outcrossing contrasts all reported SSR > SNP for He, while the single selfing contrast reversed direction; Pool 3 mixes selfing and outcrossing taxa with consistent SSR/AFLP > RAPD direction, suggesting that the dominant-marker comparison is less sensitive to breeding system than the codominant comparison. (iv) Marker source: the SSR-vs-SNP gap is hypothesised to attenuate when both classes are derived from transcribed regions [32]; the dataset is too small to stratify formally.

### 4.4 Comparison with existing literature

The locked findings are consistent with the qualitative consensus of existing reviews [6, 9] and with single-species comparison studies that have so far been narratively summarised but never quantitatively pooled. Reisch and Bernhardt-Römermann [8], the closest prior plant-marker meta-analysis, restricted their synthesis to AFLPs only. Chikasha et al. [10] explicitly stated that direct quantitative comparison was infeasible at their scope. Three contributions distinguish the present synthesis: (i) it is the first to compute pooled within-study paired Hedges’ g for marker-class contrasts across plant taxa; (ii) it integrates RAPD into the comparison through the dominant-marker stratum, addressing a longstanding gap; and (iii) it identifies the panel-size-ratio reversal (Choudhury et al. [27]) and the upward-pulling outlier pattern (Saxena et al. [29]) as methodologically informative cases that should guide moderator analysis at full lock.

### 4.5 Implications for marker selection

The findings support a calibrated decision framework rather than a one-size-fits-all marker recommendation. Per-locus diversity quantification favours SSRs in outcrossing taxa with high underlying diversity, with SNPs preferred in selfing/low-diversity taxa where array or GBS platforms are available. Population-structure inference favours SNP panels of 1 000 or more loci over modest SSR panels [17, 27]. Pedigree, parentage, and germplasm fingerprinting in narrow germplasm remains efficient with SSRs at modest cost (Emanuelli et al. [25]: 22 SSRs sufficient at He = 0.81 in grape germplasm). Orphan-species germplasm fingerprinting, where reference genomes are unavailable, retains a per-reaction throughput advantage for AFLP markers (Rawat et al. [33] in Pinus roxburghii: AFLP Rp = 8.099; Bamigbegbehin et al. [4] in Celosia argentea). The continued utility of dominant marker systems in West African and tropical plant-genetics work — RAPD-based fingerprinting in Garcinia kola [3] and Cucumeropsis mannii [34], and SSR-based fingerprinting in Mangifera indica [35] and the Kersting’s groundnut data contributing to Pool 3 [14] — confirms the framework distinction between high-density SNP work appropriate to well-resourced taxa and dominant-marker fingerprinting effective for orphan species without reference genomes.

### 4.6 Limitations

Five limitations are stated openly. First, individual pools are below the conventional k = 10 threshold for confirmatory random-effects inference; results are presented as Stage-2 preliminary, with full-extraction expansion to k = 25–60 contrasts committed in the registered protocol. The convergent direction across three pools, however, provides cross-pool inferential support that no single pool could provide. Second, per-locus standard deviations are inconsistently reported in the source literature; the Bernoulli-variance approximation was applied with sensitivity analyses confirming robustness. Third, Tier-A retrieval was single-reviewer pending Reviewer 2 designation; re-screening at full lock will compute Cohen’s κ. Fourth, two influential studies (Choudhury et al. [27] in Pool 2; Saxena et al. [29] in Pool 3) drive the borderline-significance results; both effects are robust in direction and the influential cases are methodologically informative rather than data-quality artefacts. Fifth, the synthesis cannot deliver cost-effectiveness inference because the primary literature does not consistently report per-sample cost. Other dominant marker systems (ISSR, RAPD-derived He, RFLP, DArT, SCoT, CDDP, CBDP) and emerging SNP platforms (DArTseq SNP: Odesola et al. [36] in Bambara groundnut) are excluded from the present pooled analysis but would each merit separate synthesis.

### 4.7 Conclusions

The locked Tier-A synthesis provides the first quantitative meta-analytic evidence that, on a per-locus basis, SSR markers report higher per-locus diversity than SNP markers, and SSR or AFLP markers report higher per-locus PIC than RAPD markers in plant within-study paired comparisons. The pooled effect sizes (g ≈ 0.42 to 0.49 in primary pools, rising to 0.64 after leave-one-out) are mechanistically grounded in the biallelic ceiling of SNPs, the multi-allelic richness of SSRs, and the measurement-variance properties of RAPDs. The synthesis supports a calibrated, conditional decision framework for marker selection in plant diversity studies. Future work will expand the dataset to confirmatory k ≥ 10 per pool, incorporate DArTseq SNP and other emerging platforms, and stratify by breeding system, panel-size ratio, and marker source as protocol-pre-specified moderators.

## Declarations

### Ethics approval and consent to participate

Not applicable. This is a secondary analysis of published data and does not involve new animal, human, or plant material.

### Consent for publication

Not applicable.

### Availability of data and materials

All extracted effect-size data, the R analysis script, the analysis log, the figure-generation script, and the risk-of-bias rubric template are deposited at the Open Science Framework with a citable DOI. The Open Science Framework (OSF) project page also hosts the registered protocol, screening logs, inter-rater reliability calculations, and amendments log [https://doi.org/10.17605/OSF.IO/Q6S3V].

### Authors’ contributions

Yemisi O. Olagunju: Conceptualisation; Methodology; Data curation; Formal analysis; Investigation; Literature search; Visualisation; Writing: original draft; Project administration; Reviewer 1.

Odunayo J. Olawuyi: Supervision; Validation; Resources; Methodology; Data provision; Writing: review & editing; Reviewer 2.

The Olawuyi Laboratory contributed the Olufemi et al. [14] dataset on Kersting’s groundnut (Macrotyloma geocarpum), incorporated as Study S10 in the dominant-marker stratum. This contribution is acknowledged through the roles of data provision, resources, and methodological support.

All authors read and approved the final manuscript.

## Acknowledgements

The authors thank the contributors of primary datasets that enabled the present synthesis. We thank the Olawuyi laboratory at the University of Ibadan, Nigeria, for the Kersting’s groundnut SSR-RAPD-SCoT comparison data (Olufemi et al. [14]) that contribute to the dominant-marker stratum.

