## Supplementary material for "Diversity Assessment with SNP, SSR, AFLP, and RAPD Markers in Plants: A Systematic Review and Meta-Analysis": https://osf.io/kpz8g/files/hndcb (from another version of this preprint: version 1, 2026-07-06)

**Supplementary Materials
Comparative Effectiveness of SNP, SSR, AFLP, and RAPD Markers in Plant Genetic Diversity Assessment**

*Companion file to BMC Plant Biology submission (v2 with Amendment 2)*

### **Supplementary Table S1. Stage-1 + Amendment 2 systematic search execution log**

Consensus.app peer-reviewed corpus, executed 6 May 2026 (Stage-1) and 7 May 2026 (Amendment 2 RAPD-targeted searches).

*Stage-1 searches (6 May 2026):*

| **Query** | **Yield** | **Eligible (prima facie)** |
| --- | --- | --- |
| SNP versus SSR markers genetic diversity comparison PIC heterozygosity | 19 | 12 |
| Meta-analysis molecular markers genetic diversity (filter: meta-analysis) | 20 | 3 |
| SNP SSR AFLP cereals wheat rice maize sorghum diversity | 20 | 8 |
| SNP SSR AFLP forest trees fruit trees olive grape apple | 20 | 6 |
| SNP SSR AFLP marker efficiency multiplex resolving discriminating | 20 | 5 |
| SSR microsatellite SNP ornamental rose chrysanthemum lily orchid | 20 | 2 |
| SSR SNP comparison legumes chickpea common bean cowpea soybean | 20 | 4 |

*Amendment 2 supplementary searches (7 May 2026):*

| **Query** | **Yield** | **Eligible (prima facie)** |
| --- | --- | --- |
| RAPD SSR comparison genetic diversity plants polymorphism information content | 20 | 10 |
| RAPD AFLP comparison genetic diversity plants paired analysis | 20 | 8 |
| RAPD SNP marker comparison genetic diversity plant cultivars | 20 | 2 |

Combined: ~140 unique candidate records after deduplication; ~50 prima-facie eligible at title/abstract; 15 confirmed at locked Tier-A extraction.

### **Supplementary Table S2. Pool 3 — Dominant-marker stratum (Amendment 2)**

Six within-study paired contrasts of codominant or AFLP markers vs RAPD on per-locus PIC.

| **ID** | **Author (year)** | **Taxon** | **Comparison** | **n acc.** | **Marker A \| RAPD loci** | **PIC A \| PIC RAPD** | **Hedges' g [95% CI]** |
| --- | --- | --- | --- | --- | --- | --- | --- |
| S10 | Olufemi 2024 | Macrotyloma geocarpum | SSR vs RAPD | 20 | 5 \| 5 | 0.304 \| 0.192 | 0.262 [−0.629, 1.152] |
| S11 | Hussein 2023 | Mangifera indica | SSR vs RAPD | 17 | 30 \| 30 | 0.735 \| 0.378 | 0.749 [ 0.344, 1.155] |
| S13 | Dar 2017 | Sesamum indicum | SSR vs RAPD | 47 | 18 \| 22 | 0.194 \| 0.186 | 0.020 [−0.421, 0.460] |
| S14 | Saxena 2024 | Pterocarpus santalinus | SSR vs RAPD | 40 | 6 \| 20 | 0.840 \| 0.204 | 1.474 [ 0.541, 2.406] |
| S15 | Joshi 2020 | Eleusine coracana | SSR vs RAPD | 40 | 5 \| 9 | 0.370 \| 0.314 | 0.097 [−0.677, 0.871] |
| S19 | Radwan 2021 | Zea mexicana | AFLP vs RAPD | 16 | 5 combos \| 14 | 0.210 \| 0.150 | 0.134 [−0.591, 0.859] |
| **Pool 3a (k=6)** | **—** | **—** | **REML, KH** | **—** | **—** | **—** | **0.419 [−0.121, 0.960]** |
| **Pool 3b (k=5; SSR-RAPD only)** | **—** | **—** | **REML, KH** | **—** | **—** | **—** | **0.477 [−0.218, 1.172]** |

Heterogeneity Pool 3a: τ² = 0.130; I² = 56.5%; Q[5] = 12.0, p = 0.035; 95% prediction interval [−0.65, 1.49]. Heterogeneity Pool 3b: τ² = 0.175; I² = 64.5%; Q[4] = 11.4, p = 0.023; 95% PI [−0.88, 1.83].

### **Supplementary Table S3. Pool 3b leave-one-out sensitivity**

SSR-vs-RAPD sub-stratum (k = 5).

| **Study removed** | **Pooled g** | **95% CI** | **p** | **I² (%)** |
| --- | --- | --- | --- | --- |
| −Olufemi 2024 | 0.528 | [−0.476, 1.532] | 0.193 | 75.3 |
| −Hussein 2023 | 0.392 | [−0.637, 1.422] | 0.312 | 63.9 |
| −Dar 2017 | 0.637 | [−0.243, 1.517] | 0.105 | 48.6 |
| −Saxena 2024 | 0.320 | [−0.266, 0.906] | 0.181 | 51.8 |
| −Joshi 2020 | 0.568 | [−0.394, 1.530] | 0.157 | 72.4 |

Saxena 2024 (red sanders, study-level g = 1.47) is the most influential study; its removal reduces the pooled estimate to g = 0.320 without reversing direction. Unlike the Choudhury 2023 case in Pool 2, the influential study here is upward-pulling, not direction-reversing — Pool 3 is robust in direction, sensitive in magnitude.

### **Supplementary Note S1. Why RAPD-derived He is not pooled with codominant He**

RAPD markers, like AFLPs, are dominant: heterozygotes cannot be distinguished from dominant homozygotes by band presence. Per-locus expected heterozygosity for RAPDs must therefore be estimated under Hardy-Weinberg equilibrium assumption from band frequencies (e.g., the Lynch and Milligan 1994 correction or a Bayesian estimator), not measured directly as in codominant SSR or SNP markers.

In the Olufemi et al. (2024) study contributing study S10 to Pool 3, the authors used the simple 1−Σpᵢ² formula for all three marker classes (SSR, SCoT, RAPD), which is the codominant formula. This produces a downward-biased RAPD He estimate (0.230 in their data) that is not directly comparable with codominant SSR He. We therefore extract only PIC values for Pool 3 (per-locus information content under the codominant calculation, which the authors applied uniformly), and exclude RAPD-derived He from the synthesis. The same exclusion principle was applied at protocol design to AFLP-derived He.

This is a methodologically conservative choice. Future syntheses with re-extraction under the Lynch-Milligan correction would permit a separate dominant-marker He pool — this is identified as future work in the protocol.

### **Supplementary Table S4. Sensitivity to assumed within-study correlation r (Pool 2)**

| **r** | **Pooled g** | **95% CI** | **p** | **τ²** | **I² (%)** |
| --- | --- | --- | --- | --- | --- |
| 0.00 | 0.365 | [−0.102, 0.832] | 0.101 | 0.146 | 86.1 |
| 0.25 | 0.381 | [−0.093, 0.855] | 0.094 | 0.161 | 90.1 |
| 0.50 | 0.399 | [−0.081, 0.879] | 0.086 | 0.178 | 93.8 |
| 0.75 | 0.420 | [−0.065, 0.906] | 0.077 | 0.196 | 97.1 |

Computed on the original Pool 2 (k = 6) prior to amendment 1; values from the v2 analysis log. Re-computation on the corrected k = 5 pool will be added at full lock. The pooled estimate is robust in direction and magnitude across r ∈ [0, 0.75].

### **Supplementary Table S5. PRISMA 2020 compliance checklist (updated)**

All 27 items addressed in main manuscript or this supplementary file.

| **Item #** | **Item** | **Where addressed** | **Status** |
| --- | --- | --- | --- |
| 1 | Title | Title page | **PASS** |
| 2 | Abstract | Structured abstract | **PASS** |
| 3 | Rationale | Section 1 (Background) | **PASS** |
| 4 | Objectives | Section 1 final paragraph; H1–H4 | **PASS** |
| 5 | Eligibility criteria | Section 2.1; OSF Section F | **PASS** |
| 6 | Information sources | Section 2.1; full list at OSF | **PASS** |
| 7 | Search strategy | Suppl Note S1 in v1 / Appendix 1 OSF | **PASS** |
| 8 | Selection process | Section 2.1; Reviewer 2 designation pending | **PARTIAL** |
| 9 | Data collection process | Section 2.2 | **PASS** |
| 10a–b | Data items | Section 2.2; OSF Appendix 3 | **PASS** |
| 11 | Study risk of bias | Section 2.3; OSF Appendix 2 | **PASS (rubric); per-study at lock** |
| 12 | Effect measures | Section 2.2 | **PASS** |
| 13a–f | Synthesis methods | Section 2.3 | **PASS** |
| 14 | Reporting bias | Section 3.3 | **PASS (acknowledged underpowered)** |
| 15 | Certainty assessment | Section 4.6 | **PARTIAL — GRADE at lock** |
| 16a | Study selection results | Figure 1 (PRISMA flow) | **PASS** |
| 16b | Study characteristics | Section 3.1; Table 1 (k=15) | **PASS** |
| 17 | Risk of bias in studies | Section 4.6; rubric in OSF | **PARTIAL — at lock** |
| 18 | Results of individual studies | Tables 2, 3 | **PASS** |
| 19 | Results of syntheses | Section 3.5; Table 4 | **PASS** |
| 20 | Reporting biases | Section 3.3 | **PARTIAL** |
| 21 | Certainty of evidence | Section 4.6 | **PARTIAL** |
| 22 | Discussion — interpretation | Sections 4.1–4.4 | **PASS** |
| 23 | Discussion — limitations | Section 4.6 | **PASS** |
| 24a–c | Registration / protocol / amendments | Declarations; OSF; amendments log Section 2.4 | **PASS** |
| 25 | Support / funding | Declarations | **PARTIAL — USER INPUT** |
| 26 | Competing interests | Declarations | **PARTIAL — USER INPUT** |
| 27 | Availability of data, code, materials | Declarations | **PASS** |
